# Protection of nascent DNA at stalled replication forks is mediated by phosphorylation of RIF1 intrinsically disordered region

**DOI:** 10.1101/2021.11.04.467328

**Authors:** S. Balasubramanian, M. Andreani, J.G. Andrade, T. Saha, J. Garzón, W. Zhang, A. Rahjouei, D.B. Rosen, B.T. Chait, A.D. Donaldson, M. Di Virgilio

## Abstract

RIF1 is a multifunctional protein that plays key roles in the regulation of DNA processing. During repair of DNA double-strand breaks (DSBs), RIF1 functions in the 53BP1-Shieldin pathway that inhibits resection of DNA ends to modulate the cellular decision on which repair pathway to engage. Under conditions of replication stress, RIF1 protects nascent DNA at stalled replication forks from degradation by the DNA2 nuclease. How these RIF1 activities are regulated at the post-translational level has not yet been elucidated. Here, we identified a cluster of conserved ATM/ATR consensus SQ motifs within the intrinsically disordered region (IDR) of mouse RIF1 that are phosphorylated in proliferating B lymphocytes. We found that phosphorylation of the conserved IDR SQ cluster is dispensable for the inhibition of DSB resection by RIF1, but is essential to counteract DNA2-dependent degradation of nascent DNA at stalled replication forks. Therefore, our study identifies a key molecular switch that enables the genome-protective function of RIF1 during DNA replication stress.

## Introduction

Control of DNA processing is a crucial determinant for the preservation of genome stability during both DNA repair and DNA replication. In the context of DNA double-strand break (DSB) repair, nucleolytic processing of DNA ends acts as a key defining step in the regulation of repair pathway choice (Chapman et al., 2012; Scully et al., 2019). Extensive 5’ to 3’ resection of DSBs inhibits repair by nonhomologous end joining (NHEJ) but is a pre-requisite for homology-dependent repair processes (homologous recombination, HR, and alternative end joining, A-EJ) (Chang et al., 2017; Symington, 2016). These pathways are differentially engaged to mediate physiological DSB repair according to the cellular context, cell cycle phase, and type of break (Chang et al., 2017; Chapman et al., 2012; Scully et al., 2019). As a result, dysregulated DSB end processing can lead to unproductive or aberrant repair reactions with dramatic consequences at both cellular and systemic levels, as evidenced during repair of programmed DSBs in B lymphocytes undergoing class switch recombination (CSR) and of stochastic DNA replication-associated breaks in BRCA1-mutated cells.

Immunoglobulin (Ig) CSR is the process occurring in mature B lymphocytes that enables the formation of different Ig classes or isotypes, thus diversifying the effector component of immune responses (Methot and di Noia, 2017). At the molecular level, CSR is mediated by a deletional recombination reaction at the Ig heavy chain locus (*Igh*) that replaces the constant (C) gene for the basal IgM isotype with one of the downstream C genes encoding a different Ig class (Yewdell and Chaudhuri, 2017). The reaction is initiated by the formations of multiple programmed DSBs at internally repetitive DNA stretches, known as switch (S) regions, preceding the recombining C regions (Saha et al., 2021). Productive CSR events occur *via* protection of DSBs from nucleolytic resection, which enables NHEJ-mediated inter-S-region repair (Boboila et al., 2012; Saha et al., 2021). Defects in DSB end protection lead to unscheduled processing of S region breaks, which combined with the close break proximity and the repetitive nature of these DNA stretches, favors local, hence unproductive, intra-S-region recombination reactions, and results in immunodeficiency (Boersma et al., 2015; Chapman et al., 2013; Dev et al., 2018; di Virgilio et al., 2013; Escribano-Díaz et al., 2013; Findlay et al., 2018; Ghezraoui et al., 2018; Gupta et al., 2018; Hakim et al., 2012; Ling et al., 2020; Noordermeer et al., 2018; Panchakshari et al., 2018; Reina-San-Martin et al., 2007; Xu et al., 2015; Yamane et al., 2013).

Conversely, extensive processing is essential for repair of DNA replication-associated breaks, which employ HR as the physiological repair pathway (Scully et al., 2019; Symington, 2016). In this context, the HR protein BRCA1 specifically counteracts DSB end protection, thus enabling resection and HR (Bunting et al., 2010; Tarsounas and Sung, 2020). Absence of BRCA1 causes persistent protection of DNA replication-associated DSBs, which interferes with their physiological repair by HR (Bunting et al., 2010). As a result, cells accumulate unrepaired DSBs and aberrant NHEJ-mediated chromosome fusions know as radials (Bouwman et al., 2010; Bunting et al., 2010). The increased levels of genome instability are responsible for the lethality of BRCA1-mutated cells and mouse models (Tarsounas and Sung, 2020). Defects in DSB end protection can relieve the inhibitory brake on resection in BRCA1-mutated cells, and partially rescues HR, genome stability and viability (Bouwman et al., 2010; Bunting et al., 2010; Cao et al., 2009; Chapman et al., 2013; Dev et al., 2018; Escribano-Díaz et al., 2013; Feng et al., 2013; Findlay et al., 2018; Ghezraoui et al., 2018; Gupta et al., 2018; Noordermeer et al., 2018; Xu et al., 2015; Zimmermann et al., 2013).

Recently, pathways that counteract the nucleolytic degradation of nascent DNA at replication forks have proven to be crucial to maintain genome stability under conditions of replication stress (Pasero and Vindigni, 2017; Rickman and Smogorzewska, 2019; Schlacher et al., 2011). Replication fork reversal is the mechanism that converts a classic three-way junction fork into a four-way junction structure *via* the annealing of the newly synthesized complementary DNA strands and the re-annealing of the parental strands (Neelsen and Lopes, 2015). This process, which results in the formation of a fourth regressed arm, appears to have a stabilizing effect on forks stalled as a consequence of DNA replication stress (Liao et al., 2018). However, reversed forks can act as the entry point for various DNA nucleases, and unrestrained processing of the newly replicated DNA in the absence of protective factors leads to accumulation of DNA breaks and hyper-sensitivity to replication stress-inducing agents (Cortez, 2015; Neelsen and Lopes, 2015).

The multifunctional protein RIF1 has emerged as a key regulator of DNA processing. During repair of DSBs, RIF1 acts in the 53BP1/Shieldin-mediated cascade that inhibits resection of DNA ends (Chapman et al., 2013; di Virgilio et al., 2013; Escribano-Díaz et al., 2013; Feng et al., 2013; Zimmermann et al., 2013). As a consequence, ablation of RIF1 in mature B cells severely impairs NHEJ-repair of CSR DSBs, and leads to immunodeficiency in mouse models (Chapman et al., 2013; di Virgilio et al., 2013; Escribano-Díaz et al., 2013). Conversely, deletion of RIF1 in BRCA1-deficient cells partially restores resection and HR-dependent repair of DNA replication-associated breaks, and reduces genome instability and cell lethality of this genetic background (Chapman et al., 2013; Escribano-Díaz et al., 2013; Feng et al., 2013; Zimmermann et al., 2013). Furthermore, recent studies have uncovered a DNA protective role of RIF1 during replication stress (Chaudhuri et al., 2016; Garzón et al., 2019; Mukherjee et al., 2019). RIF1 is recruited to stalled DNA replication forks and protects newly synthesized DNA from processing by the DNA2 nuclease (Garzón et al., 2019; Mukherjee et al., 2019). Loss of RIF1 leads to increased degradation of nascent DNA at reversed forks, exposure of under-replicated DNA and genome instability (Chaudhuri et al., 2016; Garzón et al., 2019; Mukherjee et al., 2019).

Despite the multiple contributions of RIF1 in the regulation of DNA processing and the consequences on the preservation of genome integrity, very little is known about the post-translational control of RIF1 activities in these contexts. Furthermore, although the DSB resection inhibitory function of RIF1 has been the objective of extensive investigation, little information is available about how its DNA replication fork protective role is regulated. In this study, we report the identification of a cluster of conserved SQ motifs within mammalian RIF1 that is phosphorylated in actively proliferating B lymphocytes. Abrogation of these phosphorylation events does not affect RIF1’s ability to inhibit DSB resection but severely impairs RIF1-mediated protection of stalled DNA replication forks.

## Results

### A conserved cluster of SQ sites in RIF1 intrinsically disordered region is phosphorylated in activated B cells

RIF1 is a large protein of almost 2500 amino acids in mammalian cells (Table S1) with no known enzymatic activity. While information about RIF1 structural organization is limited, analyses of RIF1 homologs across species identified two motifs that are highly conserved from yeast to mammals: the N-terminal <u>H</u>untingtin, <u>E</u>longation factor 3, <u>A</u> subunit of protein phosphatase 2A, and <u>T</u>or1 (HEAT) repeats, and the SILK-RVxF motif, whose sequence location shifted from the N-terminus to the C-terminal end during the evolution of unicellular to multicellular organisms (Fig. 1A) (Sreesankar et al., 2012; Xu et al., 2010). In vertebrates, RIF1 also exhibits a conserved C-terminal domain with a tripartite structure (Fig. 1A) (Xu et al., 2010). The region spanning between these N- and C-terminal motifs is poorly conserved and is characterized by a high degree of intrinsic disorder (Fig. 1A). Additionally, RIF1 contains multiple serine-glutamine/threonine-glutamine (SQ/TQ) motifs, which are consensus sites for phosphorylation by the DNA damage response kinases ATM and ATR (Blackford and Jackson, 2017) (Fig. 1A and Table S2).

**Figure 1.**
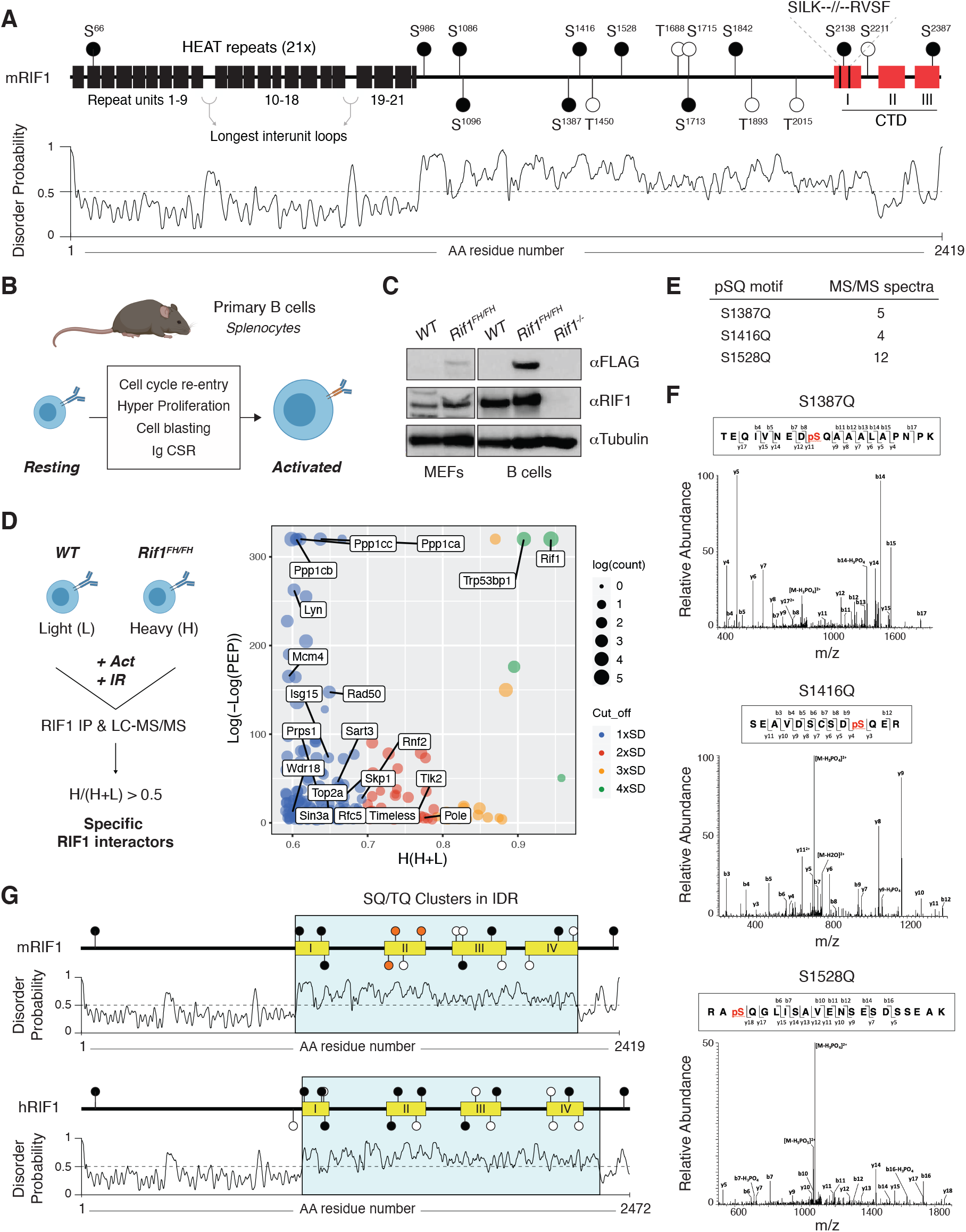
A conserved cluster of SQ motifs within RIF1 intrinsically disordered region is phosphorylated in activated B lymphocytes. (A) Top: Schematic representation of mammalian RIF1 domains and motifs. The scheme refers to the canonical sequence for mouse RIF1 (mRIF1, isoform 1, 2419 amino acids, UniProt entry Q6PR54-1). Filled and empty circle symbols represent conserved and non-conserved SQ/TQ motifs, respectively, between mRIF1 and human RIF1 (hRIF1, isoform 1, 2472 amino acids, UniProt entry Q5UIP0-1) (see also Tables S1 and S2). CTD: carboxyl-terminal domain. Bottom: Disorder profile plot of mRIF1 as determined by <u>P</u>rotein <u>D</u>is<u>O</u>rder prediction <u>S</u>ystem (PrDOS). (B) Schematic representation of key cellular changes and processes induced by the activation of mature B lymphocytes. Ig CSR: immunoglobulin class switch recombination. (C) Western blot analysis of whole cell extracts from MEFs and primary B cells derived from mice of the indicated genotypes. For each depicted antibody staining, the left and right blots represent non-contiguous portions of the same gel and film exposure. *Rif1*^*-/-*^: *Rif1*^*F/F*^*Cd19*^*Cre/+*^. (D) Left: Schematic representation of RIF1 I-DIRT in primary cultures of B cells. Light (L): light media; Heavy (H): heavy media; Act: Activation; IR: ionizing radiation; LC-MS/MS: liquid chromatography-tandem mass spectrometry. Right: Graph depicting the distribution of identified RIF1 I-DIRT proteins as a function of their H/(H+L) ratio and posterior error probability (PEP) (data from (Delgado-Benito et al., 2018)). Only proteins with posterior error probability (PEP) ≤ 10^−4^ were included in the graph. SD: standard deviation units (0.10) from the mean of the distribution (0.49); Count: number of peptides identified per protein. (E) Number of MS/MS spectra identified for the indicated phosphorylated SQ (pSQ) motif-containing peptides in different RIF1 I-DIRT preparations. (F) Representative MS/MS spectra of the RIF1 peptides encompassing phosphorylated residues S^1387^, S^1416^, and S^1528^. (G) Schematic representation of SQ/TQ motif clusters in the intrinsically disordered regions (IDRs) of mouse and human RIF1, which were defined by the PrDOS disorder profile plots (Ishida and Kinoshita, 2007). Orange filled symbols represent the conserved S^1387^, S^1416^, and S^1528^ residues identified as phosphorylated SQ motifs in mRIF1.

To identify post-translational modifications (PTMs) that modulate RIF1 functions in the maintenance of genome stability, we took advantage of the Isotopic <u>D</u>ifferentiation of Interactions as <u>R</u>andom or <u>T</u>argeted (I-DIRT) experiment that we recently performed to define RIF1 interactome in mature B lymphocytes activated to differentiate *ex vivo* (Delgado-Benito et al., 2018). In addition to experiencing programmed DSB formation and repair during Ig CSR, activated B cells undergo a proliferative burst that renders them susceptible to DNA replication stress and damage (Fig. 1B). Furthermore, activated B cells express considerably higher levels of RIF1 than their mouse embryonic fibroblasts (MEFs) counterparts (Fig. 1C).The I-DIRT approach employed primary cultures of splenocytes from mice harboring a FLAG-2xHA-tagged version of RIF1 (RIF1^FH^, (Cornacchia et al., 2012; Delgado-Benito et al., 2018)), which is expressed at physiological levels (Fig. 1C and (Delgado-Benito et al., 2018)). For the RIF1 I-DIRT experiment, activated splenocytes cultures were also irradiated, which would simultaneously increase the level and broaden the range of DNA damage-induced PTMs (Fig. 1D, (Delgado-Benito et al., 2018)). Furthermore, αFLAG-mediated pulldown of RIF1 was performed under conditions that preserved *bona fide* protein interactions and native complex formation (Fig. 1D, (Delgado-Benito et al., 2018)). The RIF1 I-DIRT experiment generated a list of high confidence interactor candidates with functions ranging from DSB repair to transcriptional regulation of gene expression (Fig. 1D and (Delgado-Benito et al., 2018)). Furthermore, a differential filtering criteria analysis uncovered an extended network of factors contributing to DNA replication initiation, elongation and fork protection (Fig. 1D). Altogether, these observations indicate that activated B cells provide an ideal model system to probe for RIF1 multiple biological functions and prompted us to re-evaluate RIF1 I-DIRT datasets for potentially relevant PTMs.

Analysis of post-translationally modified RIF1 peptides from different I-DIRT preparations revealed phosphorylation to be the predominant PTM, with the majority of phospho-residues being serines followed by either a proline or a glutamic acid (SP or SE) (data not shown). Among all SQ/TQ motifs present in mouse RIF1, S^1387^Q, S^1416^Q, and S^1528^Q were reproducibly found to be phosphorylated across independent RIF1 I-DIRT datasets (Fig. 1, E and F). These SQ motifs exhibit a relatively high degree of conservation across species (Table S1 and S2). More interestingly, S^1387^Q, S^1416^Q, and S^1528^Q (S^1403^Q, S^1431^Q, and S^1542^Q in hRIF1) are located in close proximity to each other and form a defined cluster of SQ sites in the intrinsically disordered region (IDR) of both mouse and human RIF1 (IDR-CII SQs) (Fig. 1G).

We concluded that in activated B lymphocytes, RIF1 is phosphorylated at a conserved cluster of SQ motifs within its IDR.

**Table S1.**
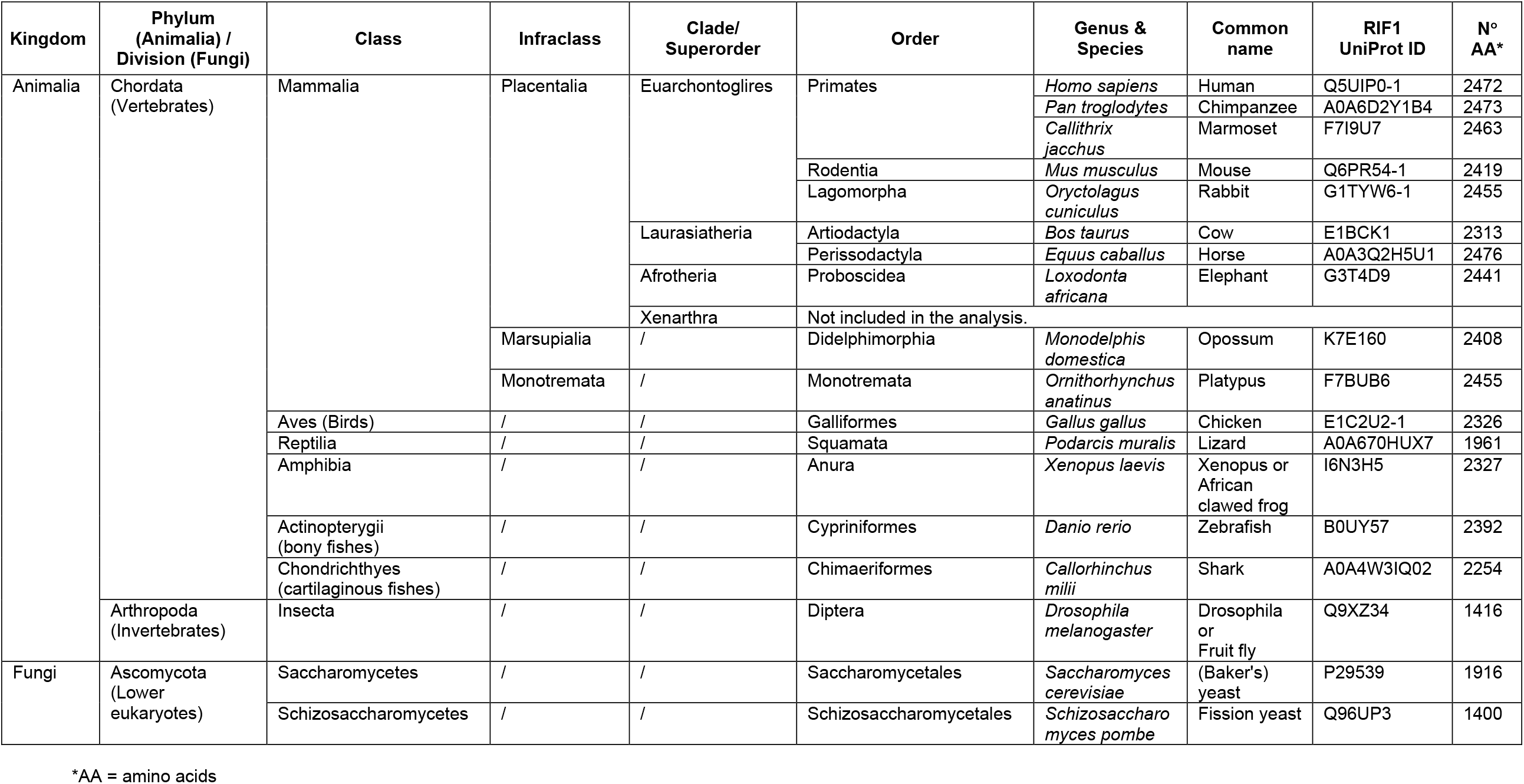
List of RIF1 protein homologs across representative species from the Animalia and Fungi kingdoms.

**Table S2.**
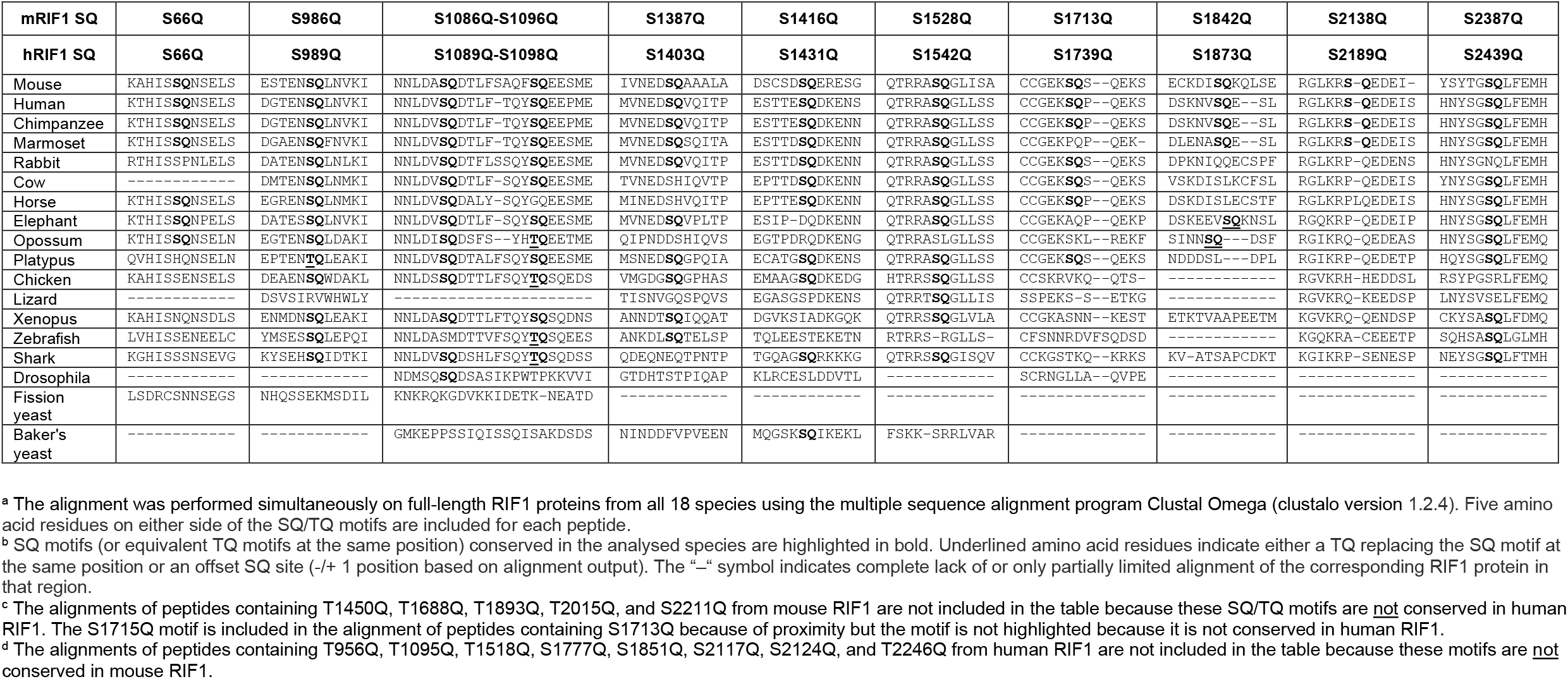
Alignment of peptides containing SQ/TQ motifs conserved between mouse and human RIF1 proteins across representative species from the Animalia and Fungi kingdoms.

## A genetic engineering-amenable B cell model system for the assessment of DSB end protection outcomes

Phosphorylation of residues within IDRs has been reported to affect protein functions in a variety of cellular contexts (Bah and Forman-Kay, 2016; Wright and Dyson, 2015). Therefore, we decided to assess the contribution of the IDR-CII SQ phosphorylation to the regulation of RIF1 activities in DNA repair. RIF1 inhibits resection of DSBs downstream 53BP1 during both aberrant repair of DNA replication DSBs in the absence of BRCA1 and physiological end joining of CSR breaks in G1 in B cells (Chapman et al., 2013; di Virgilio et al., 2013; Escribano-Díaz et al., 2013; Feng et al., 2013; Zimmermann et al., 2013). Therefore, to determine if phosphorylation of the IDR-CII modulates RIF1 role in DSB end protection, we monitored both types of repair in cells expressing phospho-mutant RIF1.

To assess for aberrant (radial chromosome formation) and physiological (CSR) repair events in the same cellular context, we opted to perform our analysis in HR-deficient, yet CSR-proficient, CH12 cells bearing hypomorphic *Brca1* mutations (Fig. 2A). CH12 is a well-characterized mouse B cell lymphoma line that recapitulates the molecular mechanism and regulation of CSR (Nakamura et al., 1996). Furthermore, CH12 cells display a stable near-diploid genome that can be easily and efficiently manipulated by somatic gene targeting (Delgado-Benito et al., 2020, 2018; Sundaravinayagam et al., 2019). These features render CH12 the preferred model system over B cells isolated from the available BRCA1-mutated mouse models, which: 1) are refractory to classic transfection methods; 2) do not allow for transduction-based reconstitution studies of large proteins like RIF1; and 3) whose primary nature precludes genetic manipulation for knock-in generation.

**Figure 2.**
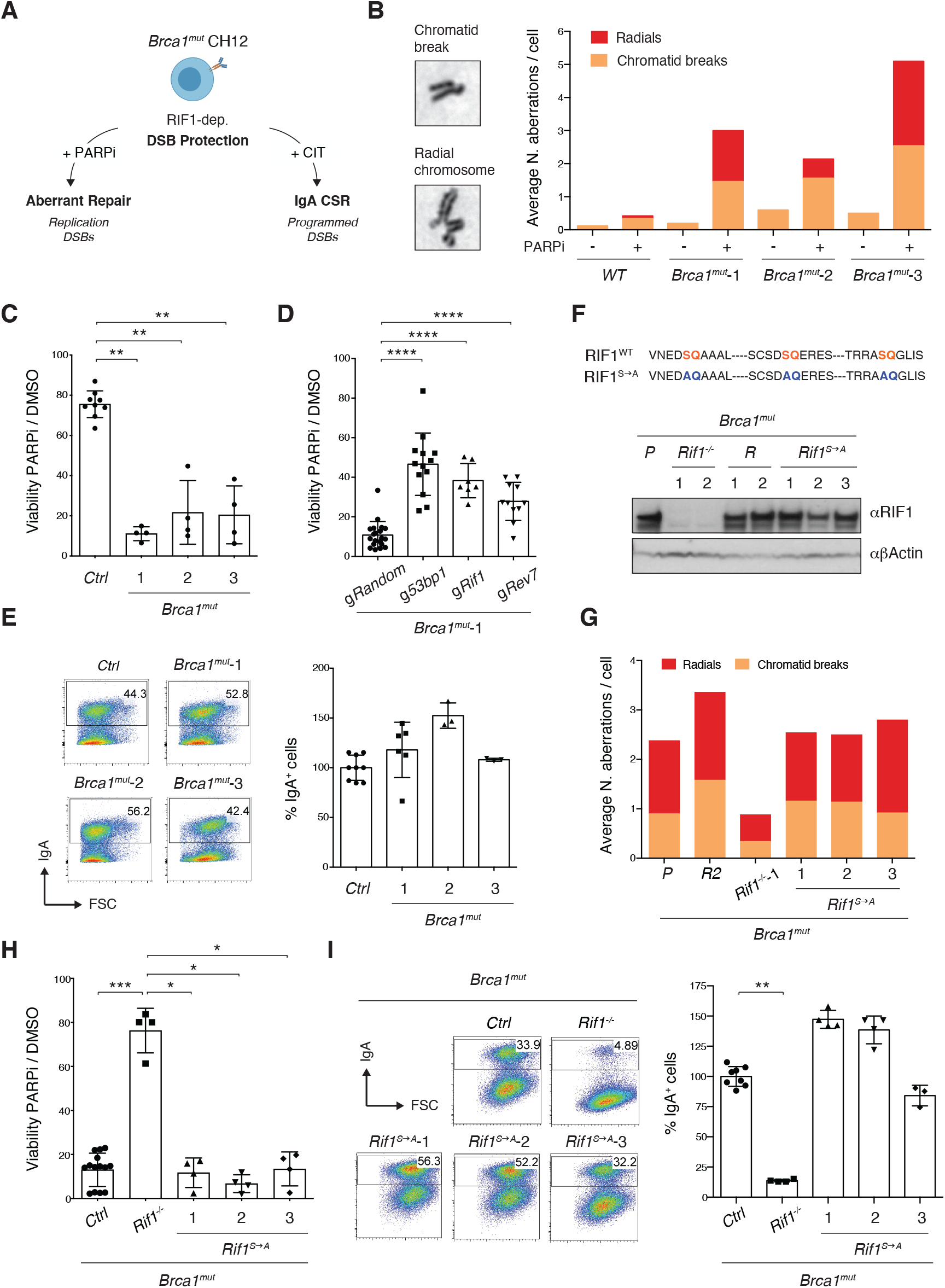
Phosphorylation of RIF1 at the conserved IDR-CII SQ motifs is dispensable for its roles in DSB end protection. (A) Schematic representation of BRCA1-deficient CH12 model system’s versatility to investigate both pathological and physiological consequences of RIF1-mediated DSB end protection. CIT: α<u>C</u>D40, IL-4 and <u>T</u>GF β B cell activation cocktail. (B) Left: Representative images of chromosomal aberrations typically associated with HR-deficiency (chromatid breaks and radial chromosomes). Right: Graph summarizing the average number of chromosomal aberrations in the parental CH12 cell line (WT sample) and selected *Brca1*^*mut*^ clonal derivatives following 1 μM PARPi treatment for 24 h (n = 50 metaphases analyzed per genotype). Breakdown of the same data into actual number of aberrations per cell is shown in Fig. S1C. (C) Residual viability of *Brca1*^*mut*^ CH12 cell lines after treatment with 1 μM of PARPi for 72 h. Graph summarizes four independent experiments per *Brca1*^*mut*^clonal derivative. Residual viability was calculated as percentage of cell viability of PARPi-over DMSO-treated cultures. The control (*Ctrl*) samples comprise parental WT CH12 cells and clonal cell lines generated by targeting CH12 cells with gRNAs against random sequences not present in the mouse genome (validated Random clones, *Brca1*^mut^*R*). (D) Residual viability of *Brca1*^*mut*^-1 CH12 cells nucleofected with random gRNAs (*Random*), or *53bp1, Rif1*, and *Rev7*, and treated for 72 h with 1 μM of PARPi. Graph summarizes eleven independent experiments. (E) Left: Representative flow cytometry plots measuring CSR to IgA in activated cell lines of the indicated genotype. Right: Summary graphs for at least three independent experiments per *Brca1*^*mut*^ cell line, with CSR% levels within each experiment normalized to the average of controls (parental WT CH12 and one Random clone), which was set to 100. (F) Top: Amino acid sequence in the IDR-CII SQ region of WT and S→A-mutated RIF1 protein. Bottom: Western blot analysis of whole cell extracts from independent cells lines of the indicated genotypes (*Rif1*^*-/-*^, control Random *R*, and *Rif1*^*S→A*^, all generated on the parental (*P*) *Brca1*^*mut*^-1 cell line background, henceforth indicated as *Brca1*^*mut*^). (G) Graph summarizing the average number of chromosomal aberrations in cells lines of the indicated genotypes following 1 μM PARPi treatment for 24 h (n = 50 metaphases analyzed per genotype). Control samples include the parental *Brca1*^*mut*^-1 cell line (*P*) and a derivative *Brca1*^mut^*R* clone. (H) Residual viability of *Brca1*^*mut*^*Rif1*^*S→A*^ cell lines after treatment with 1 μM of PARPi for 72 h. Graph summarizes four independent experiments per *Brca1*^*mut*^*Rif1*^*S→A*^ clonal derivative. The control (*Ctrl*) samples comprise parental *Brca1*^*mut*^-1 cells and *Brca1*^mut^*R* clones. (I) Left: Representative flow cytometry plots measuring CSR to IgA in activated cell lines of the indicated genotype. Right: Summary graphs for four independent experiments, with CSR% levels within each experiment normalized to the average of controls (parental *Brca1*^*mut*^-1 and one Random clone), which was set to 100. Significance in panels C, D, H, and I was calculated with the Mann–Whitney U test, and error bars represent SD. ^*^ = p ≤ 0.05; ^**^ = p ≤ 0.01; ^***^ = p ≤ 0.001; ^****^ = p < 0.0001.

To generate BRCA1-mutated CH12 cells able to support CSR, we introduced in-frame deletions specifically within exon 11 of the *Brca1* gene (Björkman et al., 2015; Bunting et al., 2010; Callen et al., 2013). Targeted deletion of *Brca1* exon 11 in mice results in the expression of a splice variant (BRCA1-Δ11) that preserves intact N-terminal RING finger domain and C-terminal BRCT repeats but lacks key motifs that are essential for BRCA1 functions (Evers and Jonkers, 2006; Xu et al., 2001, 1999). BRCA1-Δ11-expressing B cells exhibit genome instability because of impaired HR but undergo CSR as proficiently as WT cells (Bunting et al., 2010; Callen et al., 2013). We employed two different nickase gRNA pairs directed towards the 5’ region of the exon (Fig. S1A). All analyzed clones bore in-frame deletions, which are indicative of internally deleted, hypomorphic BRCA1 mutants (*Brca1*^*mut*^, Fig. S1B and data not shown). To functionally confirm the partial loss of BRCA1 function, we assessed the levels of chromosomal aberrations in response to treatment with the PARP inhibitor Olaparib (PARPi). PARPi increases the load of DNA replication breaks, and in BRCA1-deficient backgrounds it triggers the accumulation of chromatid breaks and radial chromosomes (Farmer et al., 2005). These aberrations are caused by the inability to engage physiological repair by HR, in part because of suppressed DSB end resection (Bouwman et al., 2010; Bunting et al., 2010). The resulting genome instability is responsible for the increased cell lethality associated with PARPi treatment in this genetic background (Farmer et al., 2005; Rottenberg et al., 2008). Analysis of metaphase spreads revealed that in contrast to control cells, PARPi-treated *Brca1*^*mut*^ CH12 cell lines accumulated chromatid breaks and radials with high frequency (Fig. 2B and S1C). Accordingly, all *Brca1*^*mut*^ cell lines displayed reduced viability in the presence of PARPi compared to their wild-type counterparts (Fig. 2C). We concluded that *Brca1*^*mut*^ CH12 cells exhibit genome instability-driven cell death following PARPi.

Deletion of DSB end protection factors in BRCA1-deficient cells releases the inhibition on DNA end resection, and partially rescues HR, genome stability, and, as a consequence, viability (Bouwman et al., 2010; Bunting et al., 2010; Cao et al., 2009; Chapman et al., 2013; Dev et al., 2018; Escribano-Díaz et al., 2013; Feng et al., 2013; Findlay et al., 2018; Ghezraoui et al., 2018; Gupta et al., 2018; Noordermeer et al., 2018; Xu et al., 2015; Zimmermann et al., 2013). Therefore, we monitored the consequences of ablating key components of the DSB end protection cascade in *Brca1*^*mut*^ CH12 cell lines. In bulk targeting of RIF1 as well as of the up- and downstream pathway components 53BP1 and REV7, respectively, led to a significant rescue of viability in *Brca1*^*mut*^ cells (Fig. 2D). Furthermore, *Brca1*^*mut*^*Rif1*^*-/-*^ clonal derivatives exhibited reduced levels of chromosomal aberrations and a marked increase in viability after PARPi treatment compared to *Brca1*^*mut*^ cells (Fig. S2, A to E). We concluded that *Brca1*^*mut*^CH12 cell lines recapitulate the RIF1-dependent genome instability and cell lethality typical of BRCA1-deficient backgrounds.

CH12 cells can be induced to undergo CSR to IgA with high efficiency after activation with α CD40, IL-4 and TGF β (CIT cocktail, (Nakamura et al., 1996)). NHEJ repair of CSR breaks in CH12 mimics the molecular requirements of the physiological process in primary B cells. Accordingly, *Brca1*^*mut*^ CH12 cell lines were able to undergo stimulation-dependent CSR to levels comparable to WT CH12 cells (Fig 2E), whereas deletion of RIF1 in these cells dramatically impaired CSR (Fig. S2F).

Altogether, these findings show that *Brca1*^*mut*^ CH12 cell lines allow for the investigation of both outcomes of RIF1-mediated DSB end protection: aberrant repair of DNA replication-associated DSBs and physiological end joining of CSR breaks.

**Figure S1.**
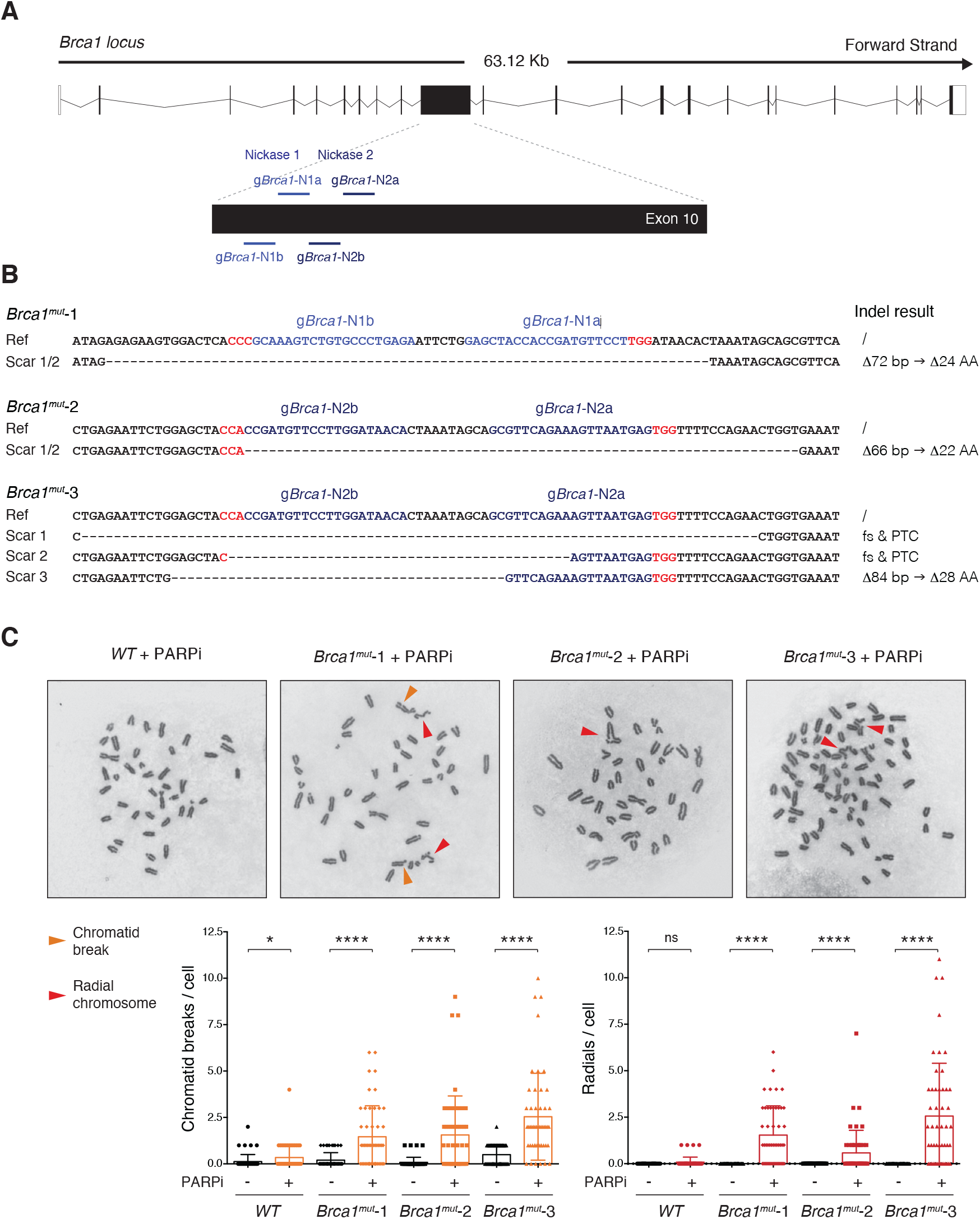
BRCA1-mutated CH12 cell lines recapitulate the genomic instability of BRCA1 deficiency. (A) Scheme of mouse *Brca1* genomic locus (scheme adapted from Ensembl ENSMUST00000017290.10) and location of gRNAs used in this study. The nickase gRNA pairs are g*Brca1*-N1a and g*Brca1*-N1b for Nickase 1, and g*Brca1*-N2a and g*Brca1*-N2b for Nickase 2. Please note that the targeted exon, which is the 10^th^ exon in the official Ensembl *Brca1*-201 transcript, is commonly referred to in literature (and in the text of this manuscript) as exon 11 because of an historical misannotation of one additional exon (Evers and Jonkers, 2006; Miki et al., 1994). (B) Genomic scar of the selected *Brca1*^*mut*^ clonal derivatives (*Brca1*^*mut*^-1, *Brca1*^*mut*^-2, *and Brca1*^*mut*^-3). The expected/potential consequences at the protein level are indicated for each scar. The gRNAs employed to generate each cell line are highlighted in shades of blue with the PAM sequence in red. Note that *Brca1*^*mut*^-1 and *Brca1*^*mut*^-2 bear the same CRISPR scar on both alleles whereas the *Brca1*^*mut*^-3 cell line has three genomic scars since it possesses a near-tetraploid chromosome set (see also panel C). Ref: reference sequence; Indel: insertion and/or deletion; Δ : base pairs (bp)/amino acid (AA) deletion; fs: amino acid frameshift; PTC: premature termination codon. (C) Analysis of genomic instability in *Brca1*^*mut*^ CH12 cell lines. Top: Representative metaphase spreads from parental CH12 cell line (WT sample) and selected *Brca1*^*mut*^ clonal derivatives following PARPi treatment (1 μM for 72 h). Orange and red arrows indicate examples of chromatid breaks and radial chromosomes, respectively. Bottom: Summary graphs for the number of chromatid breaks and radial chromosomes per metaphase/cell (n = 50 metaphases analyzed per genotype). Significance in panel C was calculated with the Mann–Whitney U test, and error bars represent SD. ns = not significant; ^*^ = p ≤ 0.05; ^****^ = p < 0.0001.

**Figure S2.**
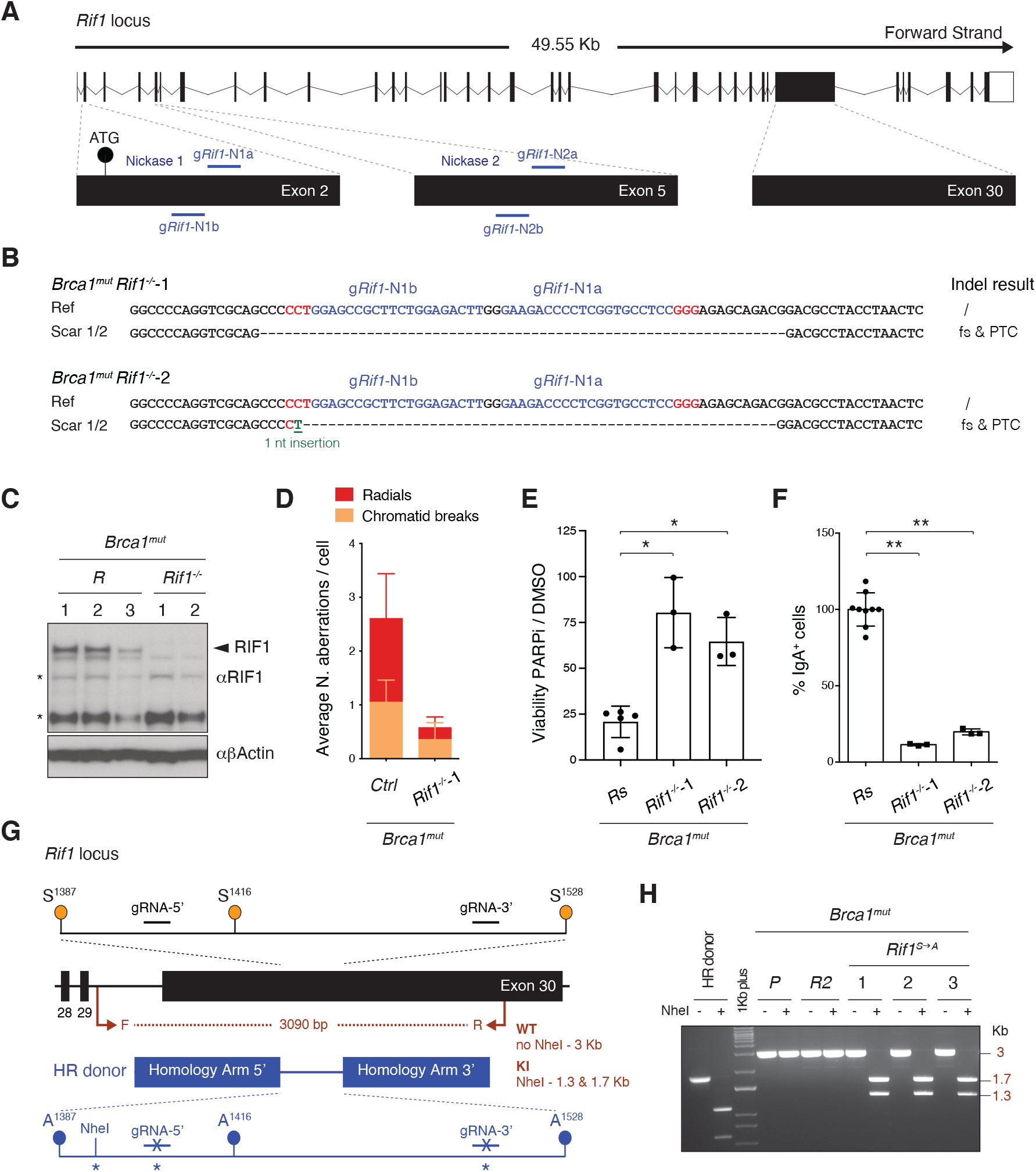
Generation of RIF1-mutant CH12 cell lines on a BRCA1-deficient background. (A) Scheme of mouse *Rif1* genomic locus (scheme adapted from Ensembl *Rif1*-201 ENSMUST00000112693.9) and location of gRNAs used in this study. The nickase gRNA pairs are g*Rif1*-N1a and g*Rif1*-N1b for Nickase 1, and g*Rif1*-N2a and g*Rif1*-N2b for Nickase 2. (B) Genomic scar of two selected/representative *Brca1*^mut^*Rif1*^*-/-*^ clonal derivatives (*Brca1*^mut^ *Rif1*^*-/-*^-1 and *Brca1*^mut^ *Rif1*^*-/-*^-2). The expected consequences at the protein level are indicated for each scar. The gRNAs employed to generate each cell line are highlighted in shades of blue with the PAM sequence in red. Ref: reference sequence; Indel: insertion and/or deletion; fs: amino acid frameshift; PTC: premature termination codon. (C) Western blot analysis of whole cell extracts from control *Brca1*^mut^*R* and *Brca1*^*mut*^*Rif1*^*-/-*^ cell lines. The arrowhead indicates RIF1 band whereas the asterisk symbol denotes aspecific bands that can be used as additional internal loading controls. (D) Graph depicting the average number of chromosomal aberrations in clonal derivatives of the indicated genotype following 1 μM PARPi treatment for 24 h (n = 50 metaphases analyzed per genotype). Graph summarizes the results of two independent experiments performed on different control cell lines (*Ctrl*, either parental *Brca1*^mut^ and/or *Brca1*^mut^*Rs*) and one *Brca1*^mut^ *Rif1*^*-/-*^ clonal derivative. (E) Residual viability of control *Brca1*^mut^*Rs* and *Brca1*^*mut*^*Rif1*^*-/-*^ cell lines after treatment with 1 μM of PARPi for 72 h. Graph summarizes three independent experiments per *Brca1*^*mut*^*Rif1*^*-/-*^ clone. (F) Dot plot depicting CSR to IgA in activated control (*Brca1*^mut^*R*) and two selected *Brca1*^mut^ *Rif1*^*-/-*^ cell lines. The graph summarizes three independent experiments per *Brca1*^mut^*Rif1*^*-/-*^cell line with CSR efficiencies within each experiment normalized to either one, or the average CSR value of two, control *Brca1*^mut^*R*, which was set to 100%. (G) Schematic representation of the knock-in strategy employed for the Ser to Ala mutagenesis at the conserved SQ motifs in the IDR-CII cluster. Location of primers (F, forward; R, reverse) and expected PCR digestion products employed for initial assessment of targeting results are indicated in dark red. KI: knock-in. “^*^” indicates silent mutations introduced to create the NheI diagnostic restriction site and to render the knocked-in sequences resistant to CRISPR-Cas9 digestion. (H) Characterization of selected *Brca1*^mut^*Rif1*^*S→A*^ CH12 cell lines by diagnostic digestion. The bands refer to the NheI-undigested and -digested products of the PCR analysis performed with the primers indicated in panel G. Note that the undigested and digested products of the donor DNA PCR fragment (digestion control) are of smaller size compared to the cell line sample fragments because they are amplified with primers inside the homology arms of the donor plasmid. The control cell line samples comprise the parental *Brca1*^*mut*^-1 cell line (P) and a *Brca1*^mut^*R* clonal derivative. Significance in panels E and F was calculated with the Mann–Whitney U test, and error bars represent SD. ^*^ = p ≤ 0.05; ^**^ = p ≤ 0.01.

### Phosphorylation of the IDR-CII SQ cluster is dispensable for RIF1 ability to inhibit DSB end resection

To investigate whether the phosphorylation status of the conserved IDR-CII SQ cluster is required for RIF1 ability to inhibit DSB end resection, we abrogated phosphorylation of S^1387^Q, S^1416^Q, and S^1528^Q motifs by serine to alanine substitutions in *Brca1*^*mut*^ CH12 cells *via* CRISPR-Cas9-mediated knock-in mutagenesis at the *Rif1* locus (Fig. 2F and S2, G and H). This knock-in approach allows the characterization of the PTM-dependent regulation of RIF1 biological functions under physiological levels of protein expression. Despite the expected HR-deficiency of *Brca1*^*mut*^ cells, we obtained several clonal derivatives that harboured the desired mutations (*Brca1*^*mut*^*Rif1*^*S→A*^), and expressed wild-type levels of RIF1 (Fig. 2F). To control for any clonality-related issue, we employed three independent clonal derivatives for all subsequent analyses.

We first asked whether preventing phosphorylation of the IDR-CII SQ cluster affected RIF1 ability to inhibit resection during repair of DNA replication DSBs. To do so, we assessed chromosomal aberrations and viability following PARPi treatment. All *Brca1*^*mut*^*Rif1*^*S→A*^ cell lines accumulated chromatid breaks and radial chromosomes to the same levels as the control *Brca1*^*mut*^ genotype (Fig. 2G) and were as sensitive to the treatment (Fig. 2H). In contrast, *Brca1*^*mut*^*Rif1*^*-/-*^cells exhibited the expected reduction in chromosomal aberrations and rescue of viability (Fig. 2, G and H). These results show that abrogation of phosphorylation events at the conserved cluster does not affect RIF1’s role in promoting genome instability in BRCA1-deficient cells.

We next assessed the contribution of IDR-CII SQ cluster phosphorylation on CSR, which is dependent on RIF1 ability to protect CSR breaks against resection (Chapman et al., 2013; di Virgilio et al., 2013; Escribano-Díaz et al., 2013). To this end, we stimulated control and *Brca1*^*mut*^*Rif1*^*S→A*^ cell lines with α CD40, TGF β and IL-4, and monitored CSR efficiencies. Whereas *Brca1*^*mut*^*Rif1*^*-/-*^ cells were, as expected, severely impaired in the process, *Brca1*^*mut*^*Rif1*^*S→A*^ cell lines all switched proficiently from expressing IgM to IgA (Fig. 2I). This finding indicates that phosphorylation of the conserved IDR-CII SQ motifs is dispensable for physiological levels of CSR.

We concluded that RIF1-mediated DSB end protection activity is not dependent on the phosphorylation of the conserved IDR-CII SQ cluster.

### Phosphorylation of the IDR-CII SQ cluster enables RIF1-dependent protection of stalled DNA replication forks

RIF1 has recently been reported to play a genome-protective role under conditions of DNA replication stress (Chaudhuri et al., 2016; Garzón et al., 2019; Mukherjee et al., 2019). RIF1 is recruited to stalled DNA replication forks where it protects nascent DNA from degradation by the DNA2 nuclease in a manner dependent on its interaction with Protein Phosphatase 1 (PP1) (Chaudhuri et al., 2016; Garzón et al., 2019; Mukherjee et al., 2019). This activity allows for timely restart of stalled forks and prevents genome instability (Garzón et al., 2019; Mukherjee et al., 2019). Given the high proliferative nature of the cellular context where phosphorylation of the conserved IDR-CII SQ motifs was originally detected (activated primary B cells, Fig. 1), we asked whether these PTMs could influence RIF1 function during replication stress.

BRCA1 plays a protective role at DNA replication forks that is independent from RIF1 (Chaudhuri et al., 2016; Garzón et al., 2019; Mukherjee et al., 2019; Schlacher et al., 2012). Therefore, to specifically address the contribution of RIF1 phosphorylation to fork protection, we first generated a set of *Rif1* knock-out and A^1387^A^1416^A^1528^-bearing phosphomutant cell lines on a BRCA1-proficient background (WT CH12 cells) (*Rif1*^*-/-*^ *and Rif1*^*S→A*^, Fig. 3A and S3, A to C). As expected, deletion of RIF1 severely impaired CSR ((Chapman et al., 2013; di Virgilio et al., 2013; Escribano-Díaz et al., 2013) and Fig. S3D), whereas, in agreement with the findings from the BRCA1-deficient background (Fig. 2I), CSR was not affected in *Rif1*^*S→A*^ cell lines (Fig. S3D).

**Figure 3.**
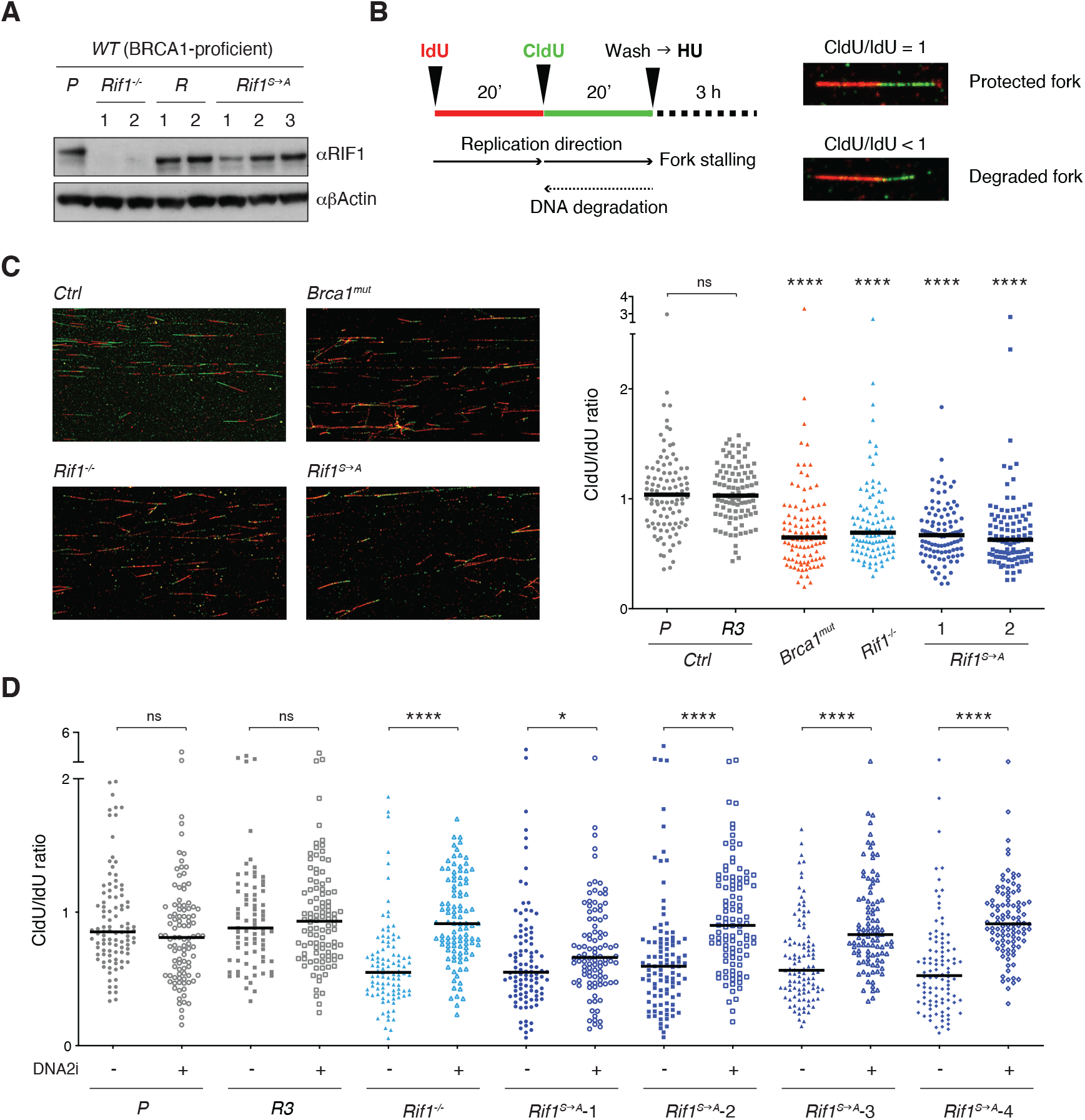
Phosphorylation of the conserved IDR-CII SQ cluster enables RIF1-dependent protection of nascent DNA at stalled replication forks. (A) Western blot analysis of whole cell extracts from independent cells lines of the indicated genotypes (*Rif1*^*-/-*^, control Random clones *R*, and *Rif1*^*S→A*^, all generated on the parental - *P* - WT CH12 background). (B) Left: Schematic representation of the DNA fiber assay employed to assess protection of nascent DNA at stalled replication forks. Right: Representative images of protected and degraded DNA fibers. (C) Left: Representative fields for the analysis of nascent DNA degradation following 3 h treatment with 4 mM HU in CH12 cells of the indicated genotypes. Right: Graph summarizing the quantification of CldU/IdU ratio for n = 100 DNA fibers analyzed per genotype (1 and 2 indicate two different *Rif1*^*S→A*^ clonal derivatives). The graph is representative of three independently performed experiments. (D). Graph summarizing the quantification of CldU/IdU ratio for n ≥ 100 DNA fibers analyzed per genotype in HU-treated cells in the absence/presence of 0.3 μM DNA2i (four different *Rif1*^*S→A*^ clonal derivatives were employed). The graph is representative of two independently performed experiments. Significance in panels C and D was calculated with the Mann–Whitney U test, and the median is indicated. Significance for each cell line in the graph in panel C was calculated in reference to the parental CH12 (*P*) sample. ns = not significant; ^*^ = p ≤ 0.05; ^****^ = p < 0.0001.

Next, we applied the DNA fiber assay to monitor the degradation of nascent DNA at forks that were stalled *via* treatment with Hydroxyurea (HU) (Fig. 3B). HU interferes with DNA synthesis by inhibiting ribonucleotide reductase, the rate-limiting enzyme in dNTP synthesis (Singh and Xu, 2016). Both *Rif1*^*-/-*^ and *Brca1*^*mut*^ genotypes exhibited the expected fork degradation phenotype (Fig. 3C), thus indicating that the protective pathways mediated by these factors are active also in CH12 cells. Interestingly, *Rif1*^*S→A*^ clonal derivatives showed increased degradation of stalled forks compared to controls, and to the same levels observed in *Rif1*^*-/-*^cells (Fig. 3C), thus suggesting that abrogation of these IDR-CII SQ phosphorylation events prevents RIF1 function at the forks.

RIF1 protective role at stalled forks is dependent on the ability of its interactor PP1 to dephosphorylate and inactivate DNA2, which in turn limits the nuclease-mediated processing of DNA replication forks (Garzón et al., 2019; Mukherjee et al., 2019). To confirm the DNA2-dependency of the fork degradation phenotype observed in cells expressing phospho-mutant RIF1 protein, we repeated the DNA fiber assay in the presence of the DNA2 inhibitor NSC-105808 (DNA2i) (Garzón et al., 2019; Kumar et al., 2017). Analogously to the result observed in *Rif1*^*-/-*^ cells (Fig. 3D and (Garzón et al., 2019; Mukherjee et al., 2019)), DNA2i treatment rescued the fork degradation phenotype in all HU-treated *Rif1*^*S→A*^ clonal derivatives (Fig. 3D).

We concluded that phosphorylation of the conserved IDR-CII SQ cluster enables RIF1-dependent inhibition of DNA2 activity and protection of nascent DNA at stalled replication forks.

**Figure S3.**
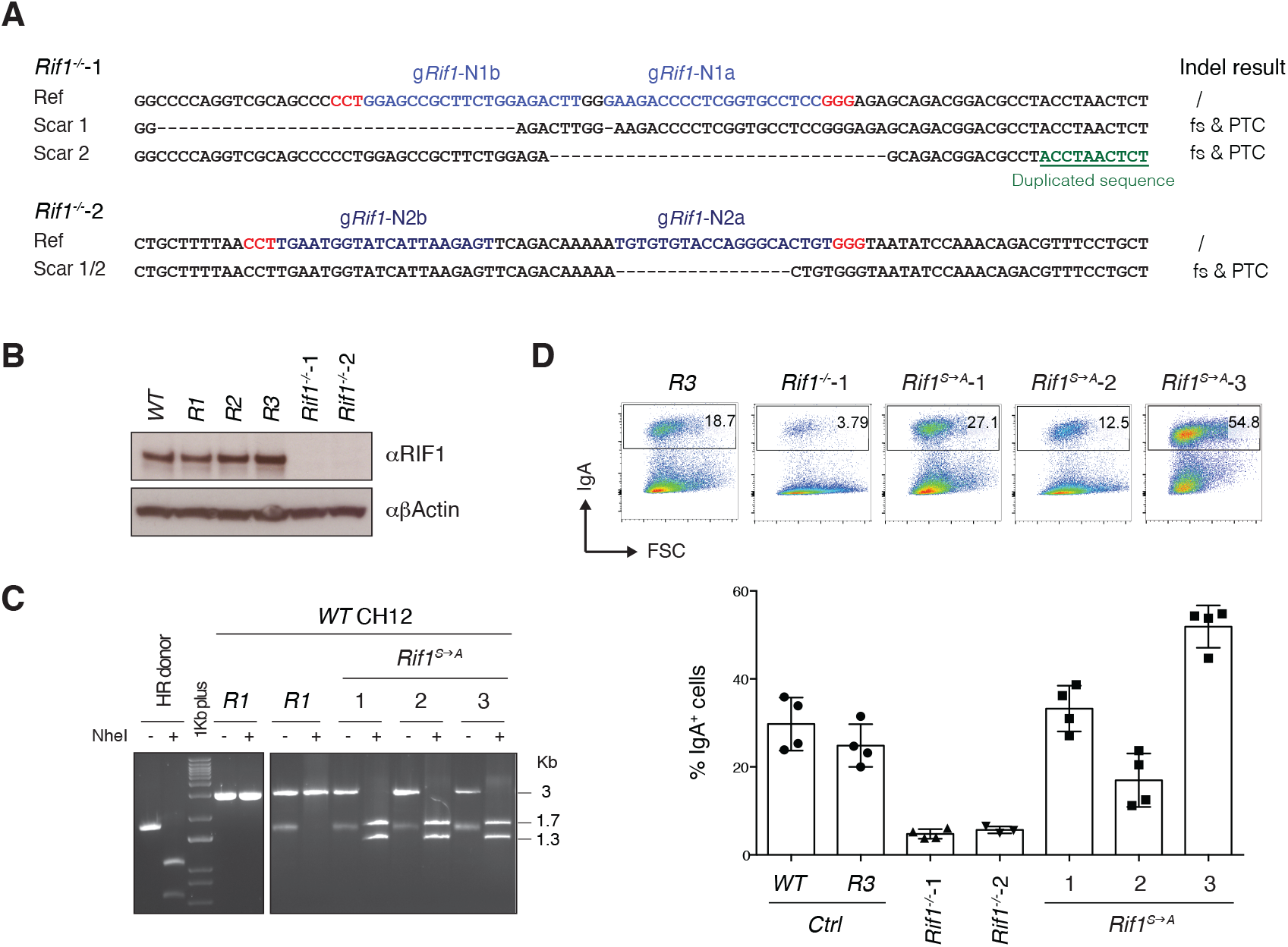
Generation of RIF1-mutant CH12 cell lines on a WT background. (A) Genomic scar of two selected *Rif1*^*-/-*^ clonal derivatives (*Rif1*^*-/-*^-1 and *Rif1*^*-/-*^-2). The expected consequences at the protein level are indicated for each scar. The gRNAs employed to generate each cell line are highlighted in shades of blue with the PAM sequence in red. Ref: reference sequence; Indel: insertion and/or deletion; fs: amino acid frameshift; PTC: premature termination codon. (B) Western blot analysis of whole cell extracts from controls and two different *Rif1* knock-out cell lines (*Rif1*^*-/-*^*-1* and *Rif1*^*-/-*^*-2*). Controls: WT parental CH12 cultures and clonal cell lines derived from targeting CH12 cells with gRNAs against random (R) sequences not present in the mouse genome (validated Random clones on the *WT* background, *R1, R2*, and *R3*). (C) Characterization of selected *Rif1*^*S→A*^ CH12 cell lines by diagnostic digestion, which followed the same scheme as for *Brca1*^mut^ *Rif1*^*S→A*^ clonal derivatives in Fig. S2, panels G and H. (D) Top: Representative flow cytometry plots measuring CSR to IgA in activated cell lines of the indicated genotype (*Rif1*^*S→A*^ -1, *Rif1*^*S→A*^ -2, and *Rif1*^*S→A*^ -3 are independent clonal derivatives). Bottom: Summary graphs for four independent experiments.

## Discussion

The regulation of DNA processing and its consequences for the preservation of genome integrity have important clinical implications. On the one hand, down-regulation or inactivating mutations in DSB end protection factors confer resistance to PARP inhibitors in BRCA1-deficient tumors in mice and a patient-derived model (Dev et al., 2018; Jaspers et al., 2013; Noordermeer et al., 2018; Xu et al., 2015). On the other hand, protection of DNA replication forks has recently been proposed as a mechanism for chemoresistance in the context of BRCA-deficiency (Chaudhuri et al., 2016). Hence, the dissection of pathways and molecular determinants in the regulation of DNA processing has profound implications for the development and improvement of targeted antitumoral treatments.

RIF1 plays at least two, and to some extent conflicting, roles in the preservation of genome integrity during DNA replication: a genome-protective role in stabilizing nascent DNA at stalled but unbroken forks, and a potentially genome-destabilizing role in regulating DNA repair by opposing resection at DSBs. Both roles depend on RIF1 ability to control DNA processing, albeit on different DNA substrates and *via* independent mechanisms: the protection of newly replicated DNA at stalled forks through PP1-induced DNA2 inactivation (Garzón et al., 2019; Mukherjee et al., 2019), and the inhibition of DSB resection at collapsed forks *via* the 53BP1-triggered cascade (Chapman et al., 2013; Escribano-Díaz et al., 2013; Feng et al., 2013; Zimmermann et al., 2013), respectively. Given the impact of these pathways on genome stability and cell viability, it is likely that multiple layers of regulations have evolved to ensure the coordination of RIF1 activities in the control of DNA processing. In this study, we identified three serine residues that are phosphorylated in hyper-proliferative B lymphocytes. This set of phospho-sites is specifically required for the role of RIF1 at stalled forks, and as such, exerts a genome protective function under conditions of DNA replication stress.

Interestingly, the identified PTMs occur within a cluster of conserved SQ motifs in the IDR of mammalian RIF1. IDRs are stretches of sequences that do not adopt any stable, defined secondary or tertiary structures (Wright and Dyson, 2015). Proteins characterized by a high degree of intrinsic disorder rapidly transition between different folding states. Phosphorylation of key residues within IDRs has been shown to influence protein folding, interaction with binding partners, and as a consequence, protein function in several biological settings (Bah and Forman-Kay, 2016; Wright and Dyson, 2015). As a relevant example, phosphorylation of 53BP1 SQ/TQ motifs within its intrinsically disordered N-terminus is essential for the DNA damage-dependent recruitment of RIF1 to sites of damage and protection against DSB resection (Chapman et al., 2013; di Virgilio et al., 2013; Escribano-Díaz et al., 2013; Feng et al., 2013; Silverman et al., 2004; Zimmermann et al., 2013).

Orthologous IDRs exhibit molecular features that are crucial for function but do not translate into any noticeable similarity at the level of primary amino acid sequences (Zarin et al., 2019, 2017). These molecular features, which include for instance length, complexity and net charge, appear to be under evolutionary selection, thus explaining how the functional output of IDRs could be maintained despite highly divergent amino acid sequences (Zarin et al., 2019, 2017). The phosphorylation of a set of IDR SQ motifs that we report in this study for mammalian RIF1 could represent such an evolutionary signature. In this regard, the IDR of *S. cerevisiae* Rif1 contains a cluster of seven SQ/TQ consensus motifs for the ATM/ATR yeast homologs Tel1/Mec1, some of which have been reported to be phosphorylated *in vivo* (Smolka et al., 2007; Sridhar et al., 2014; Wang et al., 2018). Interestingly, a recent bioRxiv manuscript showed that abrogation of phosphorylation at these seven SQ/TQ sites in yeast Rif1 impaired DNA replication fork protection after treatment with HU (Monerawela et al., 2020). Although RIF1 IDRs exhibit low conservation across evolution, the identification of a cluster of SQ/TQ motifs whose phosphorylation influences fork protection in both mammalian and yeast RIF1 hints at an evolutionary conserved mechanism, and underlying molecular feature, for the regulation of nascent DNA processing under conditions of DNA replication stress.

## Materials & Methods

### Mice and derived primary cell cultures

*Rif1*^*FH/FH*^(Cornacchia et al., 2012) and *Rif1*^*F/F*^*Cd19*^*Cre/+*^(di Virgilio et al., 2013) mice were previously described and maintained on a C57BL/6 background. Mice were kept in a specific pathogen-free (SPF) barrier facility under standardized conditions (20+/-2 °C temperature; 55%±15% humidity) on a 12 h light/12 h dark cycle. Mice of both genders were used for the experiments. All experiments were performed in compliance with the European Union (EU) directive 2010/63/EU, and in agreement with Landesamt für Gesundheit und Soziales directives (LAGeSo, Berlin, Germany).

Primary cell cultures of resting B lymphocytes were isolated from *WT, Rif1*^*FH/FH*^and *Rif1*^*F/F*^*Cd19*^*Cre/+*^ mouse spleens using anti-CD43 MicroBeads (Miltenyi Biotec), and grown in RPMI 1640 medium (Life Technologies) supplemented with 10% fetal bovine serum (FBS), 10 mM HEPES (Life Technologies), 1 mM Sodium Pyruvate (Life Technologies), 1X Antibiotic Antimycotic (Life Technologies), 2 mM L-Glutamine (Life Technologies), and 1X 2-Mercaptoethanol (Life Technologies) at 37 °C and 5% CO2 levels. Naïve B cells were activated by addition of 25 μg/ml LPS (Sigma-Aldrich), 5 ng/ml of mouse recombinant IL-4 (Sigma-Aldrich), and 0.5 μg/ml anti-CD180 (RP/14) (BD Biosciences) to the cultures upon isolation.

Primary MEFs (pMEFs) were isolated from *WT* and *Rif1*^*FH/FH*^ mice as it follows. Pregnant mice were sacrificed on day E12.5 by cervical dislocation, and embryos were removed from uterine horns and placed individually in plates containing PBS (Thermo Fisher Scientific). Brain, tail, limbs and dark red organs were removed and the remaining tissue was transferred into fresh PBS. Tissue was treated with 2 ml of Trypsin-EDTA 0.05% (Gibco) at 37°C for 15 min, and cell suspension was passed through a syringe with gauge 18 needle. Trypsin was neutralized with DMEM medium (Life Technologies) supplemented with 10% FBS, 2 mM L-Glutamine, and Penicillin-Streptomycin (Life Technologies). pMEFs from each embryo were expanded in 25 cm plates at 37°C and 5% CO2 levels to reach 80% confluency, and either used immediately for immortalization (see below) or frozen.

### Cell lines

The cell lines employed for this study are: CH12 (CH12F3, mouse, (Nakamura et al., 1996); *Rif1*^*-/-*^ CH12 (clone 1, mouse, (Delgado-Benito et al., 2018)); WT (Random clones), *Rif1*^*-/-*^ (clone 2), and *Brca1*^*mut*^ CH12 clonal derivatives (mouse, this paper), as well as RIF1 phosphomutant CH12 cell lines generated on both WT and *Brca1*^*mut*^ backgrounds (mouse, this paper); *WT* and *Rif1*^*FH/FH*^ immortalized mouse embryonic fibroblasts (iMEFs, this paper). iMEFs were generated by immortalization of the pMEFs cultures described above *via* retroviral transduction of a construct expressing the SV40 T-antigen.

CH12 cells were grown in RPMI 1640 medium supplemented with 10% FBS, 10 mM HEPES, 1 mM Sodium Pyruvate, 1X Antibiotic Antimycotic, 2 mM L-Glutamine, and 1X 2-Mercaptoethanol at 37 °C and 5% CO2 levels. iMEFs were cultured in DMEM medium supplemented with 10% FBS, 2 mM L-Glutamine, and Penicillin-Streptomycin at 37 °C and 5% CO2 levels.

### Identification of RIF1 phosphoresidues

RIF1 phosphoresidues were identified *via* analysis of RIF1 I-DIRT samples prepared from primary B cell cultures as previously described (Delgado-Benito et al., 2018), with the only difference that preparations with varying concentrations of Glutaraldehyde (1 - 5 mM) were employed. Samples were loaded on NuPAGE Bis-Tris Gels (ThermoFisher Scientific) and run for a short time to produce gel plugs. The gel samples were subjected to in-gel tryptic digestions. Peptides were extracted, purified, and analyzed by LC-MS using a Thermo Orbitrap Fusion mass spectrometer, with a Thermo Easy-nLC 1000 HPLC and a Thermo Easy-Spray electrospray source. Isotopically labelled proteins were identified by searching against a mouse protein sequence database using the GPM software (Beavis, 2006), which was set to search for tryptic peptides whose lysines and arginines were isotopically labelled and for potential phosphorylation modifications at serines, threonines and tyrosines.

### CSR assay

CH12 cells were stimulated to undergo CSR to IgA by treatment with 1-5 μg/mL α CD40 (BioLegend), 5 ng/ml TGFβ (R&D Systems) and 5 ng/ml of mouse recombinant IL-4 for 48 h. For class switching analysis, cell suspensions were stained with fluorochrome-conjugated anti-IgA (Southern Biotech) and samples were acquired on an LSRFortessa cell analyzer (BD-Biosciences).

### CRISPR-Cas9 gene targeting & generation of CH12 clonal cell lines

Targeting of *Brca1* and *Rif1* loci for generation of indel-bearing clonal derivatives was performed with 2 gRNA pairs per gene (g*Gene*-N1a and g*Gene*-N1b for Nickase 1, and g*Gene*-N2a and g*Gene*-N2b for Nickase 2) cloned into tandem U6 cassettes in a version of pX330 plasmid (pX330-U6-Chimeric_BB-CBh-hSpCas9, Addgene #42230) mutated to express Cas9^D10A^-T2A-GFP (Nickase-1/2). The Nickase-1/2 constructs were individually transfected into CH12 *via* electroporation with Neon Transfection System (Thermo Fisher Scientific).

For the generation of *Rif1*^*S→A*^*and Brca1*^*mut*^ *Rif1*^*S→A*^ cell lines, gRNAs targeting *Rif1* exon 30 (g*RNA*-5’ and g*RNA*-3’) were cloned into tandem U6 cassettes in a variant of the original pX330 plasmid (pX330-U6-Chimeric_BB-CBh-hSpCas9, Addgene #42230) modified to express Cas9^WT^-T2A-GFP (pX330-Cas9^WT^-T2A-GFP, kind gift from Van Trung Chu, MDC). CH12 cells (parental WT and *Brca1*^*mut*^-1) were co-electroporated with the g*RNA*-5’/3’-expressing construct and a circular donor plasmid carrying the synthesized knock-in template (GeneArt^®^Invitrogen™).

For the generation of both indel- and knock-in-bearing clonal derivatives, single GFP-positive cells were sorted in 96-well plates 40 h after electroporation, and allowed to grow for ca. 12 days before expansion of selected clones. Clonal cell lines were validated at the level of genomic scar (all clonal derivatives), protein level (RIF1 in *Brca1*^mut^ *Rif1*^*-/-*^, *Brca1*^*mut*^ *Rif1*^*S→A*^, *Rif1*^*-/-*^, and *Rif1*^*S→A*^), and phenotypic consequences (BRCA1-deficiency-driven genome instability and lethality in *Brca1*^mut^). Random control cell lines were generated with gRNAs against random sequences not present in the mouse genome (random gRNAs pairs-Cas9^D10A^ constructs).

For in bulk targeting of *Brca1*^*mut*^-1 cells in the rescue-of-viability assay, gRNAs against random sequences, *53bp1, Rif1*, and *Rev7* genes were cloned into the U6 cassette of pX330-Cas9^WT^-T2A-GFP. *Brca1*^*mut*^-1 cells were transfected with the Cas9-gRNAs expressing constructs *via* electroporation, sorted for GFP-positive cells after 40 h, left to recover for 72 h, and then treated with 1μM PARPi for 72 h before assessment of cell viability.

The sequences of the gRNAs and genotyping primers employed in this study are listed in Table S3.

### Western blot analysis

Western blot analysis of protein levels was performed on whole cell lysates prepared by lysis in RIPA buffer (Sigma) supplemented with 1 mM DTT (Sigma), Complete EDTA free proteinase inhibitors (Roche) and Pierce Phosphatase Inhibitor Mini Tablets (ThermoFisher). The antibodies used for WB analysis are: anti-FLAG M2 (Sigma), anti-RIF1 (di Virgilio et al., 2013), anti-Tubulin (Abcam) and anti-βActin (Sigma).

### Cell viability & Metaphase analysis

For assessment of cell viability, CH12 cells were either mock-treated (DMSO, Carl Roth) or incubated with 1 μM PARPi (Olaparib - AZD2281, Selleckchem) for 72 h. Residual viability was expressed as percentage of cell viability of PARPi-over DMSO-treated cultures.

For genomic instability analysis, exponentially growing cells were treated with DMSO or 1 μM PARPi for 24 h followed by 45 min incubation at 37°C with Colcemid (Roche). Metaphase preparation and aberration analysis were performed as it follows. Cell pellets were resuspended in 0.075M KCl at 37°C for 15 min to perform a hypotonic shock, and washed/fixed with 3:1 methanol (VWR)/glacial-acetic acid (Carl Roth) solution for 30 min at RT. Metaphase spreads were prepared by dropping fixed cells on microscope slides, which were air-dried and placed at 42°C for 1 h. Giemsa staining was performed by using KaryoMAX Giemsa Stain Solution and 1X Gurr Buffer (tablets, Gibco). Metaphases were acquired with the Automated Metaphase Finder System Metafer4 (MetaSystems).

### DNA fiber assay

Degradation of nascent DNA at stalled forks was assessed as it follows. Exponentially growing CH12 cells were sequentially pulse-labelled with 40 μM of Idoxuridin (IdU) (Sigma-Aldrich) and 400 μM of 5-Chloro-2′-deoxyuridine thymidine (CldU) (Sigma-Aldrich) for exactly 20 min each, washed once with 1X PBS, and treated with 4 mM HU for 3 h. Cells were collected and resuspended in 1X PBS at a concentration of 3.5 × 10^5^ cells/ml. 3 μl of cell suspension was diluted with 10 μl of lysis buffer (200 mM Tris-HCl pH 7.5, 50 mM EDTA, and 0.5% (w/v) SDS) on a glass slide and incubated for 2 min at RT. The slides were titled at 15°-60°, air dried and fixed with 3:1 methanol/acetic acid for 10 min. Slides were denatured with 2.5 M HCl for 80 min, washed with 1X PBS and blocked with 5% BSA (Carl Roth) in PBS for 40 min. The newly replicated CldU and IdU tracks were labelled for 1.5 h with anti-BrdU antibodies recognizing CldU (1:500, Abcam) and IdU (1:50, BD Biosciences), followed by 45 min incubation with secondary antibodies anti-mouse Alexa Fluor 488 (1:500, Invitrogen) and anti-rat Alexa Fluor 546 (1:500, Invitrogen). The incubations were performed in the dark in a humidified chamber. DNA fibers were visualized using a Carl Zeiss LSM800 confocal microscope at a 40X objective magnification, and images were analysed using ImageJ software.

Whenever indicated, the DNA2 inhibitor NSC-105808 (Kumar et al., 2017) was added at a final concentration of 0.3μM for 24 h prior to HU addition.

### Protein sequence analysis

The sequence alignment of RIF1 orthologs was performed simultaneously on the full-length proteins from all 18 species listed in Table S1 using the multiple sequence alignment program Clustal Omega (clustalo version 1.2.4, https://www.ebi.ac.uk/Tools/msa/clustalo/).

The disorder profile plots were determined by the <u>P</u>rotein <u>D</u>is<u>O</u>rder prediction <u>S</u>ystem (PrDOS) server (https://prdos.hgc.jp/cgi-bin/top.cgi) (Ishida and Kinoshita, 2007) using the template-based prediction option and with the prediction false positive rate set to 5.0%.

### Statistical analysis

The statistical significance of differences between groups/datasets was determined by the Mann–Whitney U test for all data presented in this study. Statistical details of experiments can be found in the figure legends.

**Table S3.**
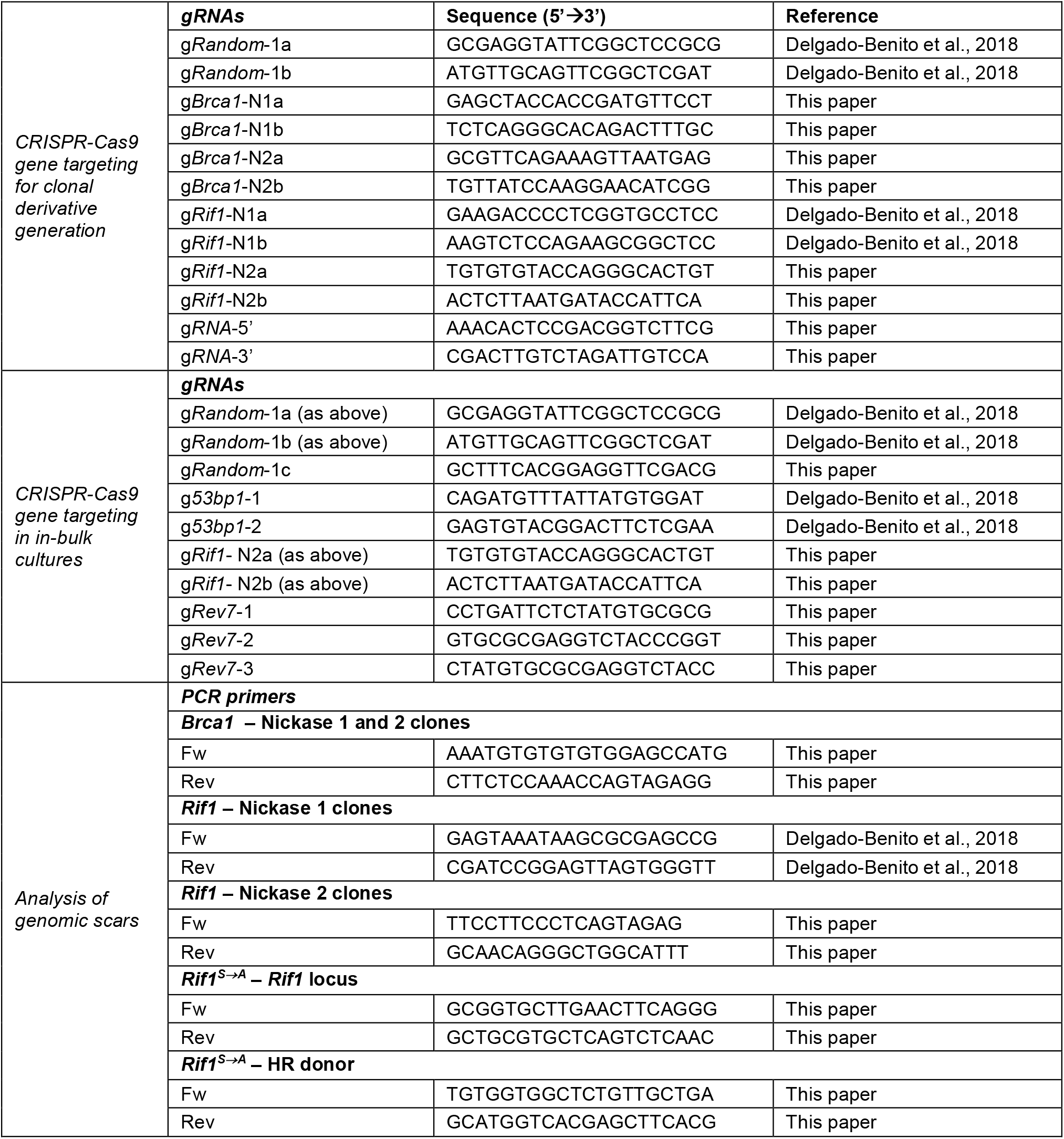
List of oligonucleotides used in this study.

## Author contributions

M.D.V., S.B., and M.A. conceived the main project idea and designed the experiments; S.B. and M.A. contributed the majority of experiments; J.G.A., T.S., J.G., A.R., and D.B.R. performed experiments; M.D.V., S.B., and M.A. analyzed and interpreted the data; W.Z. analysed the I-DIRT proteomic datasets and identified RIF1 phosphoresidues; J.G. and A.D.D. contributed their expertise and discussion on the fork protection phenotype; M.D.V., B.T.C., and A.D.D. supervised the experiments and/or analyses performed in their respective groups; M.D.V. wrote the manuscript; M.D.V. and S.B. edited the manuscript; M.D.V. obtained the main funding for the project.

## Acknowledgments

We thank all members of the Di Virgilio lab for their feedback and discussion; V. Delgado-Benito and D. Sundaravinayagam (Di Virgilio lab, MDC, Berlin) for their contributions to the project development; L. Keller (Di Virgilio lab, MDC, Berlin) for support with cloning, mutagenesis and mice genotyping; C. Brischetto (Scheidereit Lab, MDC, Berlin) for assistance with confocal microscopy; S. Hiraga (University of Aberdeen) for help with data analysis; and the MDC FACS Core Facility and Dr. HP. Rahn for support with cell sorting. This work was supported by ERC grant 638897 (to M.D.V.), the Helmholtz-Gemeinschaft Zukunftsthema “Immunology and Inflammation” ZT-0027 (to M.D.V.), and Cancer Research UK awards C1445/A19059 and DRCPGM\100013 (to A.D.D.).

## List of source data files

**Figure 1-source data 1.tiff**

Original file for the western blot analysis in Figure 1 panel C (anti-FLAG, anti-RIF1, and anti-Tubulin).

**Figure 1-source data 1 with labels.pdf**

Pdf file containing Figure 1 panel C and original scans of the relevant western blot analysis (anti-FLAG, anti-RIF1, and anti-Tubulin) with highlighted bands and sample labels.

**Figure 1-source data 2.xlsx**

Excel file containing output results of MaxQuant analysis for the potential RIF1 interactors for graph in Figure 1 panel D. Protein H/(H+L) ratios were derived using peptides’ H/L intensity values in MaxQuant output.

**Figure 2-source data 1.tif**

Original file for the western blot analysis in Figure 2 panel F (anti-RIF1).

**Figure 2-source data 2.tiff**

Original file for the western blot analysis in Figure 2 panel F (anti-βActin).

**Figure 2-source data 1 and 2 with labels.pdf**

Pdf file containing Figure 2 panel F and original scans of the relevant western blot analysis (anti-RIF1 and anti-βActin) with highlighted bands and sample labels.

**Figure 3-source data 1-right side.tif**

Original file for the western blot analysis in Figure 3 panel A (anti-RIF1 and anti-βActin).

**Figure 3-source data 1 with labels.pdf**

Pdf file containing Figure 3 panel A and original scans of the relevant western blot analysis (anti-RIF1 and anti-βActin) with highlighted bands and sample labels.

**Figure 2-figure supplement 2-source data 1.jpg**

Original file for the western blot analysis in Figure S2 panel C (anti-RIF1 and anti-βActin).

**Figure 2-figure supplement 2-source data 1 with labels.pdf**

Pdf file containing Figure S2 panel C and original scans of the relevant western blot analysis (anti-RIF1 and anti-βActin) with highlighted bands and sample labels.

**Figure 3-figure supplement 3-source data 1-left side.tif**

Original file for the western blot analysis in Figure S3 panel B (anti-RIF1 and anti-βActin).

**Figure 3-figure supplement 3-source data 1 with labels.pdf**

Pdf file containing Figure S3 panel B and original scans of the relevant western blot analysis (anti-RIF1 and anti-βActin) with highlighted bands and sample labels.

